# Low Frequency Oscillations drive EEG’s complexity changes during wakefulness and sleep

**DOI:** 10.1101/2021.12.16.472983

**Authors:** Joaquín González, Diego Mateos, Matias Cavelli, Alejandra Mondino, Claudia Pascovich, Pablo Torterolo, Nicolás Rubido

## Abstract

Recently, the sleep-wake states have been analysed using novel complexity measures, complementing the classical analysis of EEGs by frequency bands. This new approach consistently shows a decrease in EEG’s complexity during slow-wave sleep, yet it is unclear how cortical oscillations shape these complexity variations. In this work, we analyse how the frequency content of brain signals affects the complexity estimates in freely moving rats. We find that the low-frequency spectrum – including the Delta, Theta, and Sigma frequency bands – drives the complexity changes during the sleep-wake states. This happens because low-frequency oscillations emerge from neuronal population patterns, as we show by recovering the complexity variations during the sleep-wake cycle from micro, meso, and macroscopic recordings. Moreover, we find that the lower frequencies reveal synchronisation patterns across the neocortex, such as a sensory-motor decoupling that happens during REM sleep. Overall, our works shows that EEG’s low frequencies are critical in shaping the sleep-wake states’ complexity across cortical scales.

## 1. Introduction

The sleep-wake cycle is one of the most prevalent biological rhythms, being crucial to regulate physiological functions. The cycle is divided into 3 main states: wakefulness (Wake), rapid-eye movement (REM), and non rapid-eye movement sleep (NREM). At the neocortical level, Wake is characterised by asynchronous and irregular neuronal activity (Evarts, 1964; Vyazovskiy et al., 2009; Watson et al., 2016b). REM sleep is strikingly similar to Wake’s activity, with the difference that muscular activity is absent (Chase and Morales, 1983; Chase et al., 1989). In contrast, NREM exhibits neuronal synchronous silences that conform the nominative slow waves recorded in electroencephalograms (EEG) (Evarts, 1964; Vyazovskiy et al., 2009; Watson et al., 2016b; Nir et al., 2011; Todorova and Zugaro, 2019).

In order to understand cortical function during the sleep-wake cycle, classical analysis divides EEG oscillations into specific frequency bands (Buzsáki and Draguhn, 2004). These divisions stem from the oscillations being: 1) state-dependent, i.e., happen in relation to specific sleep-wake states, 2) related to different physiological functions, and 3) produced by distinct neuronal circuits. For example, frequencies up to 12 *Hz* contain the Delta (1-4 *Hz*), Theta (4-8 *Hz*) and Sigma (8-12 *Hz*) bands, which have been associated to state-dependent oscillations (Gervasoni et al., 2004; Watson et al., 2016b). On the other hand, higher frequencies, like Beta (15-30 *Hz*) or Gamma (30-150 *Hz*), have been predominantly associated to cognitive functions (Kisley and Cornwell, 2006; Kanayama et al., 2007; Bastos et al., 2015; Richter et al., 2017; Bastos et al., 2020; Wiesman et al., 2020) – even during sleep (Carr et al., 2012; Valderrama et al., 2012; Eichenlaub et al., 2020).

Recently, the classical analysis of EEG per frequency bands has been complemented by the study of EEG’s complexity (Jordan et al., 2008; Ouyang et al., 2010; Nicolaou and Georgiou, 2011; Sitt et al., 2014; Abásolo et al., 2015; Sarasso et al., 2015; Bandt, 2017; González et al., 2019, 2020; Varley et al., 2020; Hou et al., 2021; Mateos et al., 2021; Varley et al., 2021; González et al., 2021); Sarasso et al. (2021) provides a up-to-date literature review. Complexity analyses usually focus on the EEG signal as a whole, instead of its frequency components (or bands). Under this framework, it has been shown that EEG’s complexity changes according to the behavioural state, but irrespective of the animal species (including mice, rats, cats, monkeys, and humans). In particular, it has been consistently reported (Nicolaou and Georgiou, 2011; Abásolo et al., 2015; Bandt, 2017; González et al., 2019, 2020; Mateos et al., 2021; Varley et al., 2021; González et al., 2021; Pascovich et al., 2021) that Wake is a highly complex state, that complexity decreases during NREM when consciousness is lost, and that it increases during REM sleep when a state of altered consciousness emerges, i.e., dreams. However, it is still unclear how these complexity results are related to the classical frequency bands during the sleep-wake states.

Here, we study intracranial EEG (ECoG) complexity during the states of Wake, NREM, and REM sleep by dividing the ECoG recordings into low and high frequency-bands. We find that the low frequency band – including the classic Delta, Theta, and Sigma bands – contains most of the information that determines the state’s complexity. Importantly, we show that this low frequency-band preserves information across neuronal scales, from the activity of neuronal ensembles, up to the local field potentials and ECoGs. This means that our division effectively denoises ECoG signals, revealing the underlying neural oscillations. Moreover, we find novel synchronization patterns across the cortex. In particular, we find that although Wake and REM sleep have similar complexity values at the local level, cortical sensory-motor integration is severely compromised during REM sleep. Overall, our work supports classical EEG analyses that focus on the low-frequency oscillations in order to study the sleep-wake cycle, since these frequency bands contain highly relevant information.

## 2. Results

The complexity of a signal can be quantified by means of its information content, for example, by finding the signal’s Shannon Entropy (Shannon, 1948). However, for finite and real-valued signals, such as an electro-corticogram (ECoG), estimating the Shannon Entropy is challenging. Instead, we encode the ECoG signal into a finite alphabet using Ordinal Patterns (OPs), and then find the entropy – known as Permutation Entropy (Bandt and Pompe, 2002; Zanin et al., 2012). This quantification depends on the OP dimension, *D* (number of data points), and embedding delay, *τ* (resampling). In order to increase differences between close OP distributions, we use the Permutation Minimum-Entropy (PME) instead of its classical value (Zunino et al., 2015).

### 2.1. Frequency bands affect the complexity of sleep-wake states differently

Here, we study the effects that ECoG’s low and high frequency bands have on the PME of brain signals for Wake, REM, NREM sleep. We divide the recordings from 12 rats (under freely moving conditions through their sleep-wake cycle) into a frequency band ≤ 12 *Hz* and frequency band > 12 *Hz*. This division separates the classic frequencies commonly employed to visually classify sleep-wake states from the higher frequencies, which are prone to noise contamination. In particular, the low frequency-band contains different sub-bands, such as the Delta band (i.e., *δ* = {1, 4} *Hz*) related to slow-wave activity, the Theta band (i.e., θ = {4, 8} *Hz*) related to exploratory behaviour, and the Sigma band (i.e., *σ* = {8, 12} *Hz*) related to sleep spindles and memory.

Figure 1A shows the rat’s brain, where we record the activity from the primary motor cortex (M1), primary somatosensory cortex (S1) and secondary visual cortex (V2). As it can be seen from Fig. 1B, the ECoG in each cortex changes as a function of the behavioural state, from an asynchronous state during Wake and REM sleep, to a synchronous slow-wave activity during NREM sleep. We analyse the PME [Eq. (3)] per frequency band during each sleep-wake state (Fig. 1B), finding the optimal temporal scale for encoding the frequency bands into OPs. Namely, we find a state-dependent embedding delay, *τ*^⋆^, for the OP encoding of each band [Eq. (5)].

**Figure 1:**
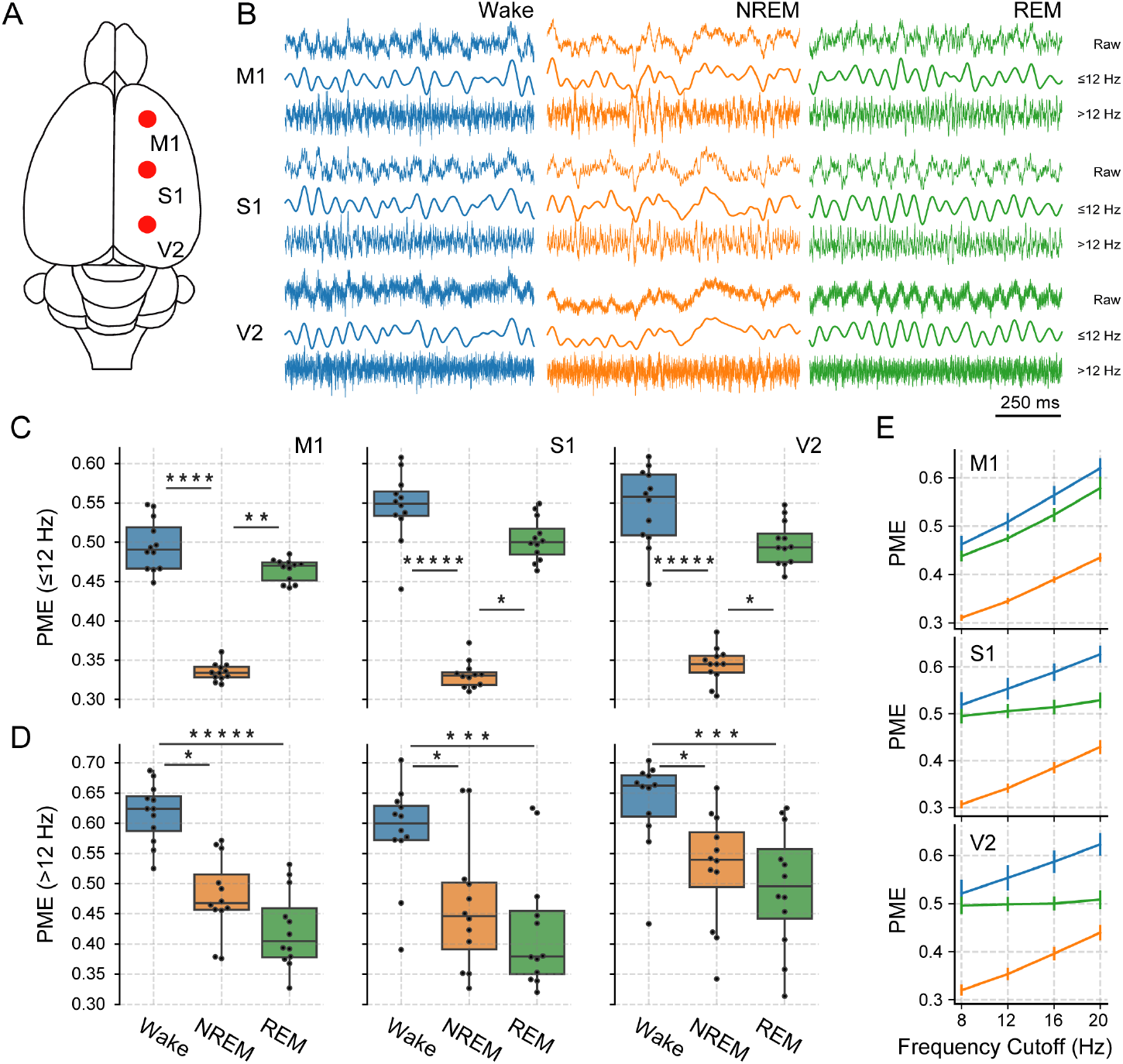
Permutation Minimum-Entropy (PME) for different sleep-wake states, cortical locations, and frequency bands. **A)** Cortical locations for the ECoG sensors: primary Motor cortex (M1), primary Somato-sensory cortex (S1), and secondary Visual cortex (V2). **B)** Examples of ECoG recordings for wakefulness (Wake), rapid-eye-movement sleep (REM), and non-REM sleep (NREM). Top traces correspond to the raw ECoG signal and bottom traces show the respective low- and high-frequency oscillations (≤ 12 *Hz* and > 12 *Hz*, respectively). Box plots show population PME values (12 rats) for the low (panel **C**) and high (panel **D**) frequency-bands at the M1, S1, and V2 cortical locations according to the sleep-wake state. PME [Eq. (3)] values obtained by encoding the ECoG signals with ordinal-patterns of dimension *D* = 5 and embedding delay, *τ*^⋆^ (Bandt and Pompe, 2002), where *τ*^⋆^ = 25 for Wake, *τ*^⋆^ = 21 for REM, and *τ*^⋆^ = 17 for NREM low frequency bands and *τ*^⋆^ = 1 for all states in the high frequency band. **E** Population average PME (error bars represent the 95% confidence interval) for each state and cortical location as a function of the maximum frequency included in the low-frequency band (frequency cut-off) for the *τ*^⋆^ in panel **C**. * *p* < 0.05, ** *p* < 0.01, **** *p* < 0.0001, ***** *p* < 0.00001

Resultant PME values for low and high-frequency bands are respectively shown in Fig. 1C and D. We can see that these values are similar across cortical areas, suggesting that PME is a cortical-area independent measurement. In particular, we find that Wake is the most complex state (regardless of the frequency band), while NREM sleep shows significantly lower complexity values for both frequency bands. Interestingly, REM’s PME strongly depends on the frequency content, showing Wake-like PME values for the lower frequencies, and NREM-like PME values for the higher frequencies.

We also show that the low-frequency band’s PME robustly differentiates the sleep-wake states even when the frequency cutoff is changed – as can be seen in Fig. 1E, where we set a range of cut-off frequencies from 8 to 20 *Hz*. For example, when the frequency cut-off is 8 *Hz*, the PME values in Fig. 1E for the frequencies ≤ 8 *Hz* are similar to those from Fig. 1C. As the cut-off is increased, higher frequencies are included in the low-frequency band, affecting the PME values and revealing a cortical dependence, where S1 and V2 behave similarly (middle and bottom panels in Fig. 1E) approaching an intermediate PME value between Wake and NREM sleep.

Overall, these results suggest that the low frequency-band contains most of the relevant information of sleep-wake states and their raw ECoG signals (i.e., before filtering). In particular, we find that in this band, Wake and REM sleep show similar PME values, which aligns with previous results by González et al. (2019) using PE. On the contrary, the high frequency-band PME variations correlate to the changes in muscular activity during sleep (Fig. S1). Consequently, for the following analysis we focus on the low-frequency band.

### 2.2. Dependence of the embedding delay on the frequency band

A signal’s information-content changes when looking at different frequency bands. This implies that the encoding needs to take into account the signal’s frequency content by adjusting the encoding parameters. In our work, when encoding an ECoG signal with ordinal patterns (OPs), we need to analyse the resultant PME [see Eq. (3)] as a function of the the embedding delay, *τ*, for each frequency band. In Fig. 2 we show the results of finding the optimal embedding delay, *τ*^⋆^ [see Eq. (5)], for the ECoG’s low frequency-band across the sleep-wake states and cortical locations.

**Figure 2:**
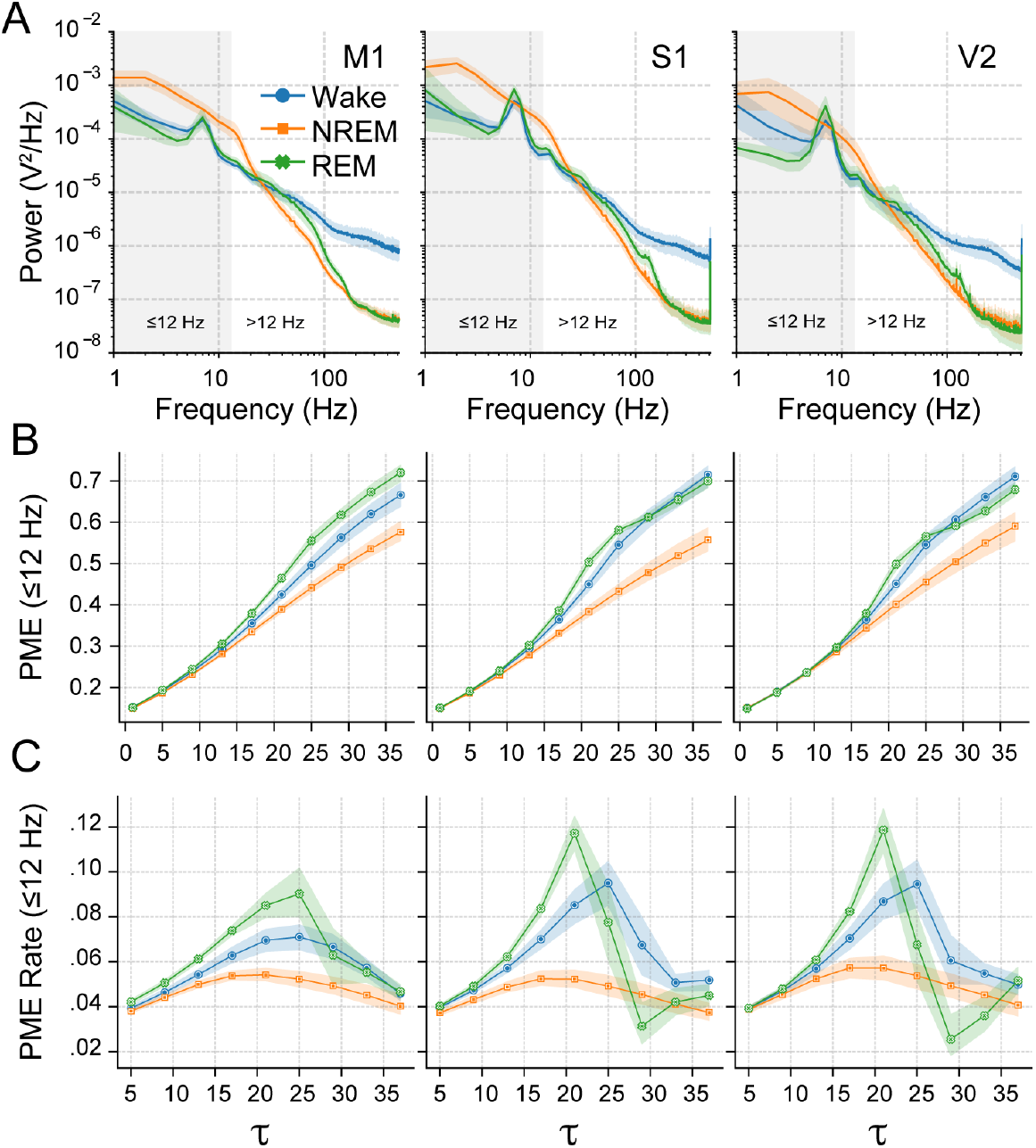
Power spectra and entropy variations for wakefulness (Wake), rapid-eye movement (REM) and non-REM (NREM) sleep. **A)** Power spectral densities for different cortical locations and sleep-wake states. Shaded rectangular areas signal the low frequency band (≤ 12 *Hz*). **B)** Permutation Minimum-Entropy (PME) values for the EEG’s low frequency-band as the embedding delay, *τ*, of the ordinal pattern (OP) is increased (i.e., data sampled at increasing steps) in each cortical location from Fig. 1A. **C)** Corresponding PME rates [Eq. (4)] from panel **B**, i.e., PME tangents as a function of *τ*. OP dimension in panels **B** and **C** is *D* = 5 data points (as in Fig. 1C). All panels show population averages (solid curves) with their 95% confidence interval (coloured shading) after 1000 bootstrap samples.

Figure 2A shows ECoG’s power spectra. We find that Wake and REM sleep have similar low-frequency content (shaded rectangle) in all neocortical areas, with a peak in the Theta range (θ = [4, 8] *Hz*). On the other hand, NREM sleep shows more power at the sleep-spindles (*σ* = [8, 12] *Hz*) and slow-wave range (*δ* = [1, 4] *Hz*). In Fig. 2B we show how the PME changes as we increase *τ* from 1 (OP constructed with consecutive data points) to 35 (OP constructed with data taken every 35 points) for the low frequency-band. We note increasing complexity values for all sleep-wake states – independently of the neocortical area. However, the growth is non-monotonic, as Fig. 2C reveals by the PME rates [see Eq. (4)]; that is, the PME tangents.

At the maximum PME rate, the encoding captures the optimal information content generated by the low frequencies. From Fig. 2C, we can see that this is obtained by an optimal embedding delay, *τ*^⋆^ [see Eq. (5)], which depends on the behavioural state (coloured curves) but is independent of the cortical location (panels). In particular, we find that Wake’s PME rate peaks at *τ*^⋆^ = 25, NREM’s at *τ*^⋆^ = 17, and REM’s at *τ*^⋆^ = 21. We note that during REM sleep, we cannot statistically differentiate between PME Rate(*τ* = 21) and PME Rate(*τ* = 25) in the M1 area (left panel in Fig. 2C), so we set *τ*^⋆^ = 21 to match the other neocortical sites. These *τ*^⋆^ values are the ones used in Fig. 1C. We then conclude that the optimal temporal scale to study ECoG dynamics solely depends on the behavioural state of the low frequencies. On the other hand, doing the same analysis to the high frequency band results in *τ*^⋆^ = 1 for all sleep-wake states and cortical locations (Fig. S2), since at this band entropy is generally driven by the high frequencies, which require a high sampling rate (i.e., that of the raw signal).

### 2.3. ECoG’s lower frequencies contain neuronal information across recording scales

Our findings show that Wake and REM sleep ECoG’s low frequency-bands have larger PME than NREM sleep. Now we analyse whether these PME values are conserved across cortical scales; that is, if Local-Field Potentials (LFP) and neuronal spiking activity has the same complexity features according to the animal’s behavioural state. In addition, we use the spiking activity binary signals to construct synthetic LFP (sLFP), which we generate by making convolutions with a decreasing exponential and then taking a population average. The resultant signal is similar to an LFP, which mainly originates from the spatial average of excitatory post-synaptic potentials (Buzsáki et al., 2012). We then perform the same analysis as in Fig. 2 to LFP and sLFP signals focusing on their low frequency-bands (≤ 12 *Hz*).

From top to bottom, Fig. 3A shows low-frequency band signals for an M1 ECoG of our experiments on 12 freely-moving rats (Cavelli et al., 2017, 2018) and a frontal cortex LFP, sLFP, and spike trains (units) of the data-set with 11 rats from the work by Watson et al. (2016b). Figure 3B shows box plots of the resultant PME values from our analysis of these recording scales in all animals, where the top panel is the same as the left panel in Fig. 1C. Here, we can see that NREM’s PMEs are significantly smaller than Wake’s and REM’s PMEs across cortical scales; that is, our findings are consistent for ECoG, LFP, and sLFPs. We also find that PME grows with increasing *τ* for all recording scales (Fig. 3C), which we previously observed in Fig. 2B for the ECoG signals. Similarly, PME rates in Fig. 3D for ECoGs, LFPs, and sLFPs exhibit comparable behaviours, where we note that *τ*^⋆^ values are always larger during Wake-fulness or REM sleep than during NREM sleep. Consequently, our findings show that low frequency-bands contain state-dependent information that stems from the spiking activity, which is conserved across the recording scales.

**Figure 3:**
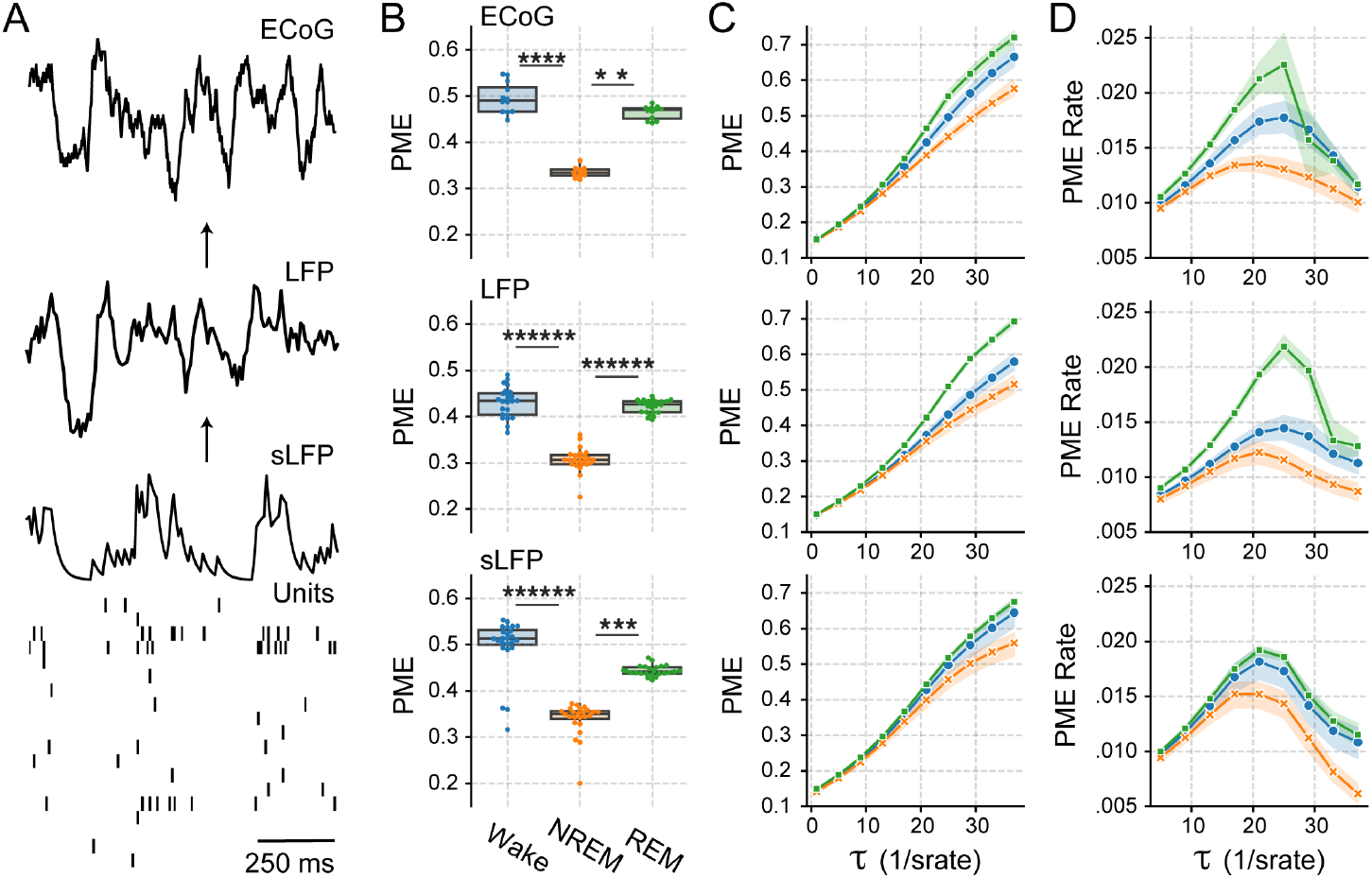
Permutation Minimum-Entropy (PME) across recording scales. **A** Brain recordings at different scales. From top to bottom, electro-corticograms (ECoG), local field potential (LFP), synthetic LFP (obtained by the convolution of the spike trains and a decreasing exponential kernel), and units (spikes from individual neurons recorded from the extracellular medium). ECoG data comes from our experiments, as in Figs. 1 and 2, but the other recordings come from the work by Watson et al. (2016b) (data-set available at: CRCNS.org). We note that we inverted both LFP and ECoG recordings for representation purposes. **B**: Box plots show PME values for the ECoGs (from Fig. 1C), LFPs, and sLFPs data-sets.* *p* < **C** PME as a function of *τ* (delay embedding); as in Fig. 2B. **C** PME Rate as a function of *τ*; as in Fig. 2**C**. The solid lines show the population average values and the shaded areas their 95% confidence interval.* *p* < 0.05, ** *p* < 0.01, **** *p* < 0.0001, ****** *p* < 0.000001

### 2.4. Sensory-motor integration is compromised during REM sleep

Having shown that the low-frequency ECoG band contains state-dependent spiking information, we now study how this activity is integrated across the neocortex. We do this by quantifying the Permutation Minimum-Mutual-Information (PMMI) between the low-frequency ECoG recordings of every pair of cortical locations, where we encode each ECoG signal into ordinal patterns (Bandt and Pompe, 2002) of length *D* = 5 and optimal embedding delay *τ*^⋆^ for each sleep-wake state (i.e., *τ*^⋆^ = 25 for Wake, *τ*^⋆^ = 21 for REM, and *τ*^⋆^ = 17 for NREM, as it can be seen from Fig. 2C).

Figure 4A shows the low-frequency ECoG signals during each sleep-wake state. During Wake (left panel in Fig. 4A), we note synchronous θ oscillations on M1 (primary motor), S1 (primary somato-sensory), and V2 (secondary visual) cortical regions. During NREM sleep (middle panel in Fig. 4A), slow waves appear almost synchronously in all cortices. On the other hand, we note that during REM sleep (right panel in Fig. 4A), the M1 cortex decouples from the rest, while S1 and V2 exhibit synchronous θ rhythms that resemble Wake.

**Figure 4:**
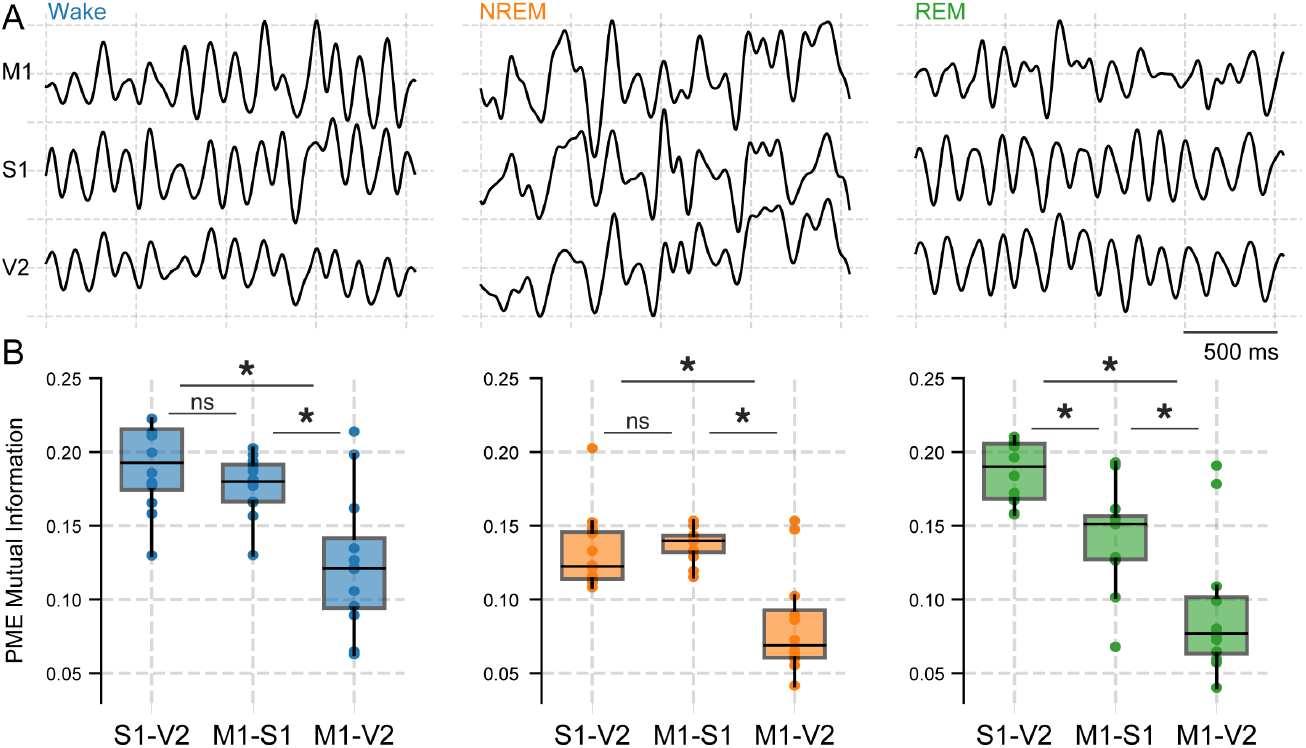
Permutation Minimum Mutual Information (PMMI) for Wakefulness, NREM and REM sleep. **A** Standardised ECoG recordings of the primary Motor cortex (M1), primary Somato-sensory cortex (S1), and secondary Visual cortex (V2) in each sleep-wake state. **B** Box-plots with the pair-wise PMMI values [Eq. (6)] between the 3 neocortical areas during each of the sleep-wake states. * signals a *P* < 0.05 Wilcoxon signed-rank test with a Benjamini-Hochberg multiple comparisons correction.

In line with these observations, PMMI values [see Eq. (6)] between cortical areas show a dependence on both the distance between cortices and the sleep-wake state (Fig. 4B). The inter-electrode distance for S1-V2 and M1-S1 is ≃ 5*mm*, but is ≃ 10*mm* for the M1-V2 combination. In particular, Fig. 4B shows that PMMI is significantly higher for cortical pairs that are ≃ 5*mm* apart in comparison to those that are ≃ 10*mm*, regardless of the sleep-wake state. However, we find a significant decrease in PMMI during REM sleep when comparing the equidistant pairs S1-V2 and M1-S1, which is absent during Wake or NREM sleep. These results point to a loss in sensory-motor integration during REM sleep that is not emerging because of cortical distances.

When comparing PMMI from different sleep-wake states, we find that REM’s M1-S1 and M1-V2 PMMI are significantly smaller than those from Wake. For example, REM’s and Wake’s population-averaged PMMI between M1 and V2 is 0.09 and 0.12 (normalised units), respectively. We highlight that this decrease in PMMI between M1 and V2 low frequencies happens even though their PME values for Wake and REM are similar (Fig. 1C). On the contrary, the decrease in PMMI values happening during NREM sleep between M1-V2 in comparison to Wake can be explained from the significantly smaller PME values that NREM shows in all cortical areas (Fig. 1C).

## 3. Discussion

Complex neural dynamics are thought to be necessary for consciousness (Tononi and Edelman, 1998; Oizumi et al., 2014). Different reports show that cortical activity exhibits complex patterns during Wakefulness, that are reduced during deep NREM sleep (Ouyang et al., 2010; Nicolaou and Georgiou, 2011; Abásolo et al., 2015; Schartner et al., 2017; Bandt, 2017; González et al., 2019, 2020; Hou et al., 2021; Mondino et al., 2021; Mateos et al., 2021) or anesthesia (Jordan et al., 2008; Sitt et al., 2014; Sarasso et al., 2015; Fagerholm et al., 2016; Thul et al., 2016; Varley et al., 2020, 2021). However, it was unclear how the different frequency bands contribute to the observed complexity changes in EEG analyses.

### 3.1. Low-frequency oscillations drive complexity changes during the states of wake and sleep

In this work, we show that the intra-cranial EEG (ECoG) frequencies up to 12 *Hz* are sufficient to reproduce the complexity variations that are typically observed across the sleep-wake states. According to our findings (Figs. 1C and 2), Delta, Theta, and Sigma bands are the most important frequencies contributing to the state’s complexity and its decrease during NREM sleep. This is supported by recent reports linking the EEG’s complexity decrease during sleep to the appearance of slow-oscillations (1-4 *Hz*) in the Delta range (Pigorini et al., 2015; D’Andola et al., 2018; Rosanova et al., 2018; Dasilva et al., 2021; Sarasso et al., 2021; González et al., 2021). In particular, we have shown that population DOWN states trap cortical dynamics into recurrent states (González et al., 2021), which explains why slow-oscillations – caused by DOWN states (Vyazovskiy et al., 2009; Nir et al., 2011; Todorova and Zugaro, 2019) – reduce the complexity of cortical activity during sleep.

Our present results also suggest that ECoG’s high frequencies (> 12 *Hz*) contain muscular tone information, which we can explain as follows. On the one hand, REM sleep shows the least complex EEG signals in the high frequency range (Fig. 1D), in spite of having neuronal activity resembling that of wakefulness (Abásolo et al., 2015; González et al., 2021). The fact that muscular tone is absent during REM sleep (Fig. S1) can explain this discrepancy between neuronal activity and ECoG’s complexity values. On the other hand, we find that the optimal delay embedding for encoding high-frequency signals is always *τ*^⋆^ = 1 (Fig. S2) for all sleep-wake states and cortical locations. This *τ*^⋆^ = 1 implies that the ordinal patterns can encode most of the information coming from frequencies in the range 1024 *Hz*/5 ∼ 200 *Hz* up to 512 *Hz* – according to Eq. (1). This range is higher than any up-to-date physiological frequency band, making the high-frequency band analysis with ordinal patterns prone to capture extra-neural sources, such as muscular tone.

Because muscular tone is intrinsically random and our approach is to maximise the entropy rates, we could be missing relevant information from the frequencies contained in the 12 to 200 *Hz* when analysing the high-frequency range. This limitation in our high-frequency band analysis could require the inclusion of an intermediate band of frequencies. Such intermediate frequencies could contain the Beta (15-30 *Hz*) and Gamma (30-150 *Hz*) bands, potentially capturing complementary information to our present work; but outside of its current scope. Nevertheless, our results remain practically unchanged when we choose a different cut-off frequency to define the low and high frequency bands (Fig. 1E), exploring cut-off values between 8 to 20 *Hz*.

### 3.2. Frequency Content of an Ordinal Pattern

When trying to measure the content of information from an ECoG signal, we need to tune the encoding to match the relevant frequencies of the signal under study. In our case, the Ordinal Pattern (OP) encoding has the embedding dimension, *D*, and delay, *τ* parameters, which set the number of points to be taken as a single OP and at which sampling rate. Consequently, depending on their values, an OP can see different frequencies, *ν*. Specifically, we can estimate the OP frequency range by

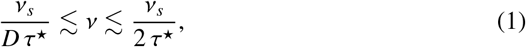

where *ν*_*s*_ is the sampling frequency of the signal (in our case, *ν*_*s*_ = 1024 *Hz*), *D* is the OP’s embedding dimension, and *τ*^⋆^ is the optimum embedding delay from Eq. (5).

Equation (1) implies that for our low frequency-band ECoG analysis (≤ 12 *Hz*), when we have *D* = 5 and *τ*^⋆^ = 25 (corresponding to wakefulness), the OP lower and upper frequency limits are approximately equal to 8 *Hz* and 20 *Hz*, respectively. This means that the *D* = 5 OP encoding will be quantifying the information content from a signal mostly within 8 and 20 *Hz*. However, we note that for different *D* and sleep-wake states, we find different *τ*^⋆^ – although independently of the cortical location. The optimal *τ*^⋆^ of the low frequency band for each sleep-wake states can be seen in Fig. 2C. On the other hand, for the high frequency-band ECoG analysis (> 12 *Hz*), we find *τ*^⋆^(*D*) = 1 independently of the embedding dimension or sleep-wake state (see Fig. S2C). In this case, the OP has an upper limit of 512 *Hz*, coinciding with the Nyquist-Shannon criterion, but a lower limit that depends on *D*, being 256 *Hz* if *D* = 4 and approximately 200 *Hz* if *D* = 5.

It is worth noting that, although Eq. (1) sets a frequency range that an OP can capture for a given *D* and *τ*^⋆^, this range only considers part of the information that an OP captures. Specifically, frequencies *ν* that are smaller than the lower bound, *ν*_*s*_/*D τ*^⋆^, are still captured by the OP. For example, a slow wave oscillation would constitute monotonically increasing or decreasing OPs, which would (strictly) have insufficient datapoints to represent the slow-wave’s period, but still contain local information about the signal and contribute to differentiate it to other frequencies.

### 3.3. Low-frequency ECoG oscillations recover neuronal dynamics

We find that we can bridge several cortical scales by focusing on the lower ECoG frequencies (Fig. 3). We note that decoding specific neuronal firing patterns from a field recording, such as an ECoG, is an ill-posed problem, but we can approximate (to a degree) the amount of information that neuronal populations generate during each sleep-wake state. In this sense, our analysis shows that Wake and REM’s neuronal dynamics and field recordings are complex across scales (Fig. 3B). In contrast, the appearance of DOWN states in neuronal populations and slow-waves in field recordings make NREM activity more predictable and less complex (González et al., 2021).

We note that although our frequency band division is a simple procedure, invariant complexity across scales disappears when considering the whole ECoG signal frequency content. In particular, if we include the high frequencies, extra-neural contamination likely confounds the complexity results and brakes the scale invariance we are finding in this work. Moreover, extra-neural sources of contamination above 20 *Hz* are already reported by Whitham (Whitham et al., 2007) for scalp EEG, which supports our decision to use a division at 12 *Hz* – making it available for scalp EEG analysis.

### 3.4. Low-frequency synchronization reveals cortical sensory-motor decoupling during REM sleep

Finally, our mutual information analysis of the low frequency band reveals particular synchronization patterns between neocortical areas during REM sleep. We find that the motor cortex decouples from sensory and visual cortices (Fig. 4), in spite of the different cortical areas maintaining complex patterns of activity. Given that the activity of these lower frequencies correlates with true neuronal dynamics (Fig. 3B), it seems unlikely that the sensory-motor decoupling is spurious. Because REM sleep is characterized by muscle atony (Chase and Morales, 1983; Chase et al., 1989), a reduction in the mutual information between the motor cortex and the rest of the brain is expected (Fig. 4B). We argue that a possible cause for this sensory-motor decoupling (i.e., less information sharing between motor areas and the rest of the neocortex), is because motor feedback signals are unable to synchronize the sensory cortices with the motor areas due to motor pathways being inhibited.

#### Experimental Procedures

*Experimental procedures* are in agreement with the National Animal Care Law (No. 18611) and with the “Guide to the care and use of laboratory animals” (8th Edition, National Academy Press, Washington DC, 2010), as well as with the approval by the Institutional Animal Care Committee, Uruguay (Exp. No 070153-000332-16). The experiments involve 12 Wistar adult rats, sustaining a controlled 12 *h* light/dark cycle (light comes on at 07:00 UYT) with unrestricted access to food and water. Veterinarians of the institution determined the animals were all in good health and we took extra care to minimise pain, discomfort, and stress in the animals. Also, we made an effort to use the minimum number of animals necessary to obtain reliable data.

*Surgical procedures* imply chronically implanting electrodes to the animals, where we follow procedures carried in previous studies by Cavelli et al. (2017, 2018). Anaesthesia is induced by a mixture of ketamine-xylazine (90 *mg*/*kg* and 5 *mg*/*kg* i.p., respectively), the rat is then positioned in a stereotactic frame, and the skull is exposed to attach 8 stainless-steel screw-electrodes, which record the intra-cranial EEG. 6 electrodes are placed bilaterally above motor (M1), somato-sensory (S1), and visual (V2) cortices. Remaining electrodes are placed in the right olfactory bulb (OB) and cerebellum (taken as the reference electrode). EMG registration is done by inserting 2 electrodes into the neck muscle. All electrodes are soldered into a 12-pin socket and fixed onto the skull with acrylic cement. At the end of the surgical procedures, an analgesic (Ketoprofen, 1 *mg*/*kg*, s.c.) is administered. After the animals recover from these surgical procedures, they are left to adapt in separate transparent cages (40×30×20 *cm*) for 1 week before data is collected. Cages contain wood-shaving material in a temperature-controlled room (set to 21-24° *C*), with water and food *ad libitum*.

*Experimental sessions* are conducted during the light period, between 10AM and 4PM UYT. Data from each rat is collected individually in a sound-attenuated recording chamber with a Faraday shield by a rotating connector that allows free movement within the cage. Polysomnographic recordings are amplified (× 1000), acquired (by a 16 bits AD converter set at a 1024 *Hz* sampling frequency), and stored using DASY Lab Software. The states of REM, NREM and Wake are determined in 10 *s* epochs. Wake is defined by low-voltage fast-waves in M1, strong theta-rhythm (4 − 7 *Hz*) in V2, and relatively high EMG activity. REM sleep is defined by low-voltage fast-frontal-waves, a regular theta-rhythm in V2, and a silent EMG (except for occasional twitches). NREM sleep is determined by the presence of high-voltage slow-cortical-waves (1 − 4 *Hz*), sleep spindles in M1, S1, and V2, and a reduction in EMG amplitudes. Additionally, a visual scoring is performed to discard artifacts and transitional states.

##### Frontal cortex data-set

We also employ the data-set from Watson et al. (2016b,a) to study population dynamics and local field potentials in the frontal cortex; freely available at CRCNS.org. For these recordings, silicon probes were implanted in frontal cortical areas of 11 male Long-Evans rats. Recording sites include the medial prefrontal cortex (mPFC), anterior cingulate cortex (ACC), pre-motor cortex/M2, and orbito-frontal cortex (OFC). Recordings took place during light hours in the home cage, including 25 sessions with mean duration of 4.8hs ± 2.2 std, at a 20 *kHz* sampling frequency. We exclude *BWRat19_032413* from our analyses because the recording lack REM sleep data. We extract Local-Field Potentials (LFPs) by applying low-pass filters to the recordings and resampling at 1250 *Hz*. We extract neuronal spikes by applying a high-pass filter at 800 *Hz* and then by detecting threshold crossings. Spike sorting is carried by means of the KlustaKwik open-source software. Sleep-wake states are identified by means of Principal Component (PC) analysis. In particular, SWS exhibits high LFP PC1 (power in the low < 20 *Hz*) and low EMG. REM sleep shows high Theta with low EMG cluster, and a diffuse cluster of low broadband LFP with high EMG. Wake has a diffuse cluster of low broadband LFP with higher EMG, and a range of Theta oscillations.

## Methods

*Pre-processing of field recordings* is done by a 1st-order Finite-duration Impulse Response (FIR1) band-pass [0.1, 450] *Hz*. We divide these signals into Low-Frequency Oscillations (LFO) by a [0.1, 12] *Hz* FIR1 band-pass and High-Frequency Oscillations (HFO) by a [13, 450] *Hz* FIR1 band-pass, which corresponds to making a division according to the classical polysomnographic frequency bands (Buzsáki and Draguhn, 2004). Then, we fix the total signal length, *T*, to a range between 3 × 10^5^ to 3 × 10^6^ data points for all cortical locations and sleep-wake states. This means that the shortest [largest] signals last *T* Δ*t* = 3 × 10^5^/1024 *Hz* ≃ 5 minutes [*T* Δ*t* ≃ 50 minutes].

*Encoding of signals into Ordinal Patterns* (OPs) [Bandt and Pompe (2002)] is done to quantify the signals’ randomness and how it changes during the sleep-wake states. This encoding involves dividing a signal, *X* = { *x*(*t*) : *t* = 0, …, (*T* − 1) Δ*t*} (where 1/Δ*t* is the sampling frequency of the signal), into sliding vectors with *D* data-points (such that *D≪T*) at a new sampling *τ*, where *D* ≥ 2 is known as the embedding dimension and *τ* as the delay embedding. For example, { *x*(*t*), *x*(*t* +*τ*), *x*(*t* +2*τ*)} is a vector at time *t* with *D* = 3 data points and sampled at *τ* ≥ 1 times. Each of these vectors is assigned an OP according to the relative magnitude of its *D* elements and how many permutations are needed to order them increasingly. In other words, an OP represents the necessary permutations needed to order the elements of the embedded vector, which has up to *D*! possible permutations. In what follows, we set *D* = 4 or 5 (meaning there are 4! = 24 or 5! = 120 possible OPs), and analyse how results change for different *τ*.

*Permutation Entropy* (Bandt and Pompe, 2002) (PE) quantifies the temporal randomness of a signal *X* after encoding it into OPs. It is defined from the probability distribution of OPs (*P*_(*D, τ*)_(*α*{*X*}), with *α* = 1, …, *D*! and *τ* the delay embedding) by

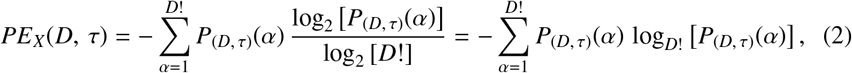

which depends on *D* and *τ* and is normalised by the maximum PE, log_2_(*D*!); namely, 0 ≤ *PE*_*X*_ ≤ 1 for any signal *X*, dimension *D*, or delay *τ*. In general, there are slight changes in the probability distribution of OPs when analysing different consciousness states. This means that PE values from Eq. (2) are similar and differences can be hindered in the statistical comparisons. In order to enhance these differences, we use the *Permutation Minimum-Entropy* (PME), which is the infinit limit of the Rényi entropy (Rényi et al., 1961; Rényi, 1965; Zanin et al., 2012; Zunino et al., 2015), is defined by

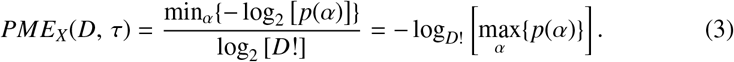

*Entropy Rates* are the incremental variations that entropy has when a parameter is changed. In our case, a PME rate is given by changes in the delay embedding, *τ* = 1, …, 40, for a fixed embedding dimension; namely, *D* = 4 or 5. We are interested on the entropy rates because of the low and high frequency-bands, which imply different relevant frequencies. Specifically, we find the PME rate of a signal *X* by

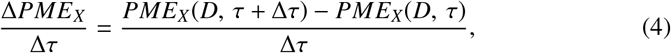

where we choose Δ*τ* = 4 for most of the PME analysis (we also explore finer values, using Δ*τ* = 1; results not shown here). In particular, we optimise *τ* by selecting the maximum PME for each cortical location and sleep-wake state. Namely,

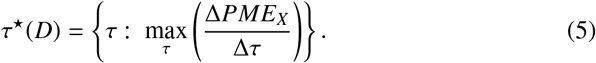

*Mutual Information, I*(*X, Y*), is the amount of shared information between 2 random signals, *X* and *Y*. It is a non-linear measure of the correlation between the signals, found from *I*(*X, Y*) = *H*(*X*) + *H*(*Y*) − *H*(*X, Y*), *H*(*X*) [*H*(*Y*)] being the marginal entropy of signal *X* [*Y*] and *H*(*X, Y*) being their joint entropy. In this work, we use the PME [Eq. (3)] to quantify the entropy of a signal, so we use this entropy to quantify a *Permutation Minimum-Mutual-Information* (PMMI) between pairs of signals. Namely,

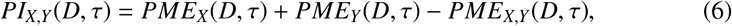

where *PME*_*X,Y*_ (*D, τ*) is the joint permutation minimum-entropy at a given *D* and *τ* (meaning that we are comparing both signals after they have been encoded into OPs).

### Statistics

We present data as regular boxplots showing the median, the 1st and 3rd quartiles, and the distribution range. Because of the complexity metrics we analyse, we employ non-parametric statistics. In particular we use the Friedman test (available with the scipy.stats) to compare the results among states (Wake-NREM-REM) with the Siegel post-hoc test applying the Benjamini-Hochberg false discovery rates correction (available with the scikitlearn python 3 package (https://scikit-learn.org/stable/)). We set *p* < 0.05 for a result to be considered significant.

## Author Contributions

- J.G. and D.M.: data processing and analysis
- M.C. and A.M. and J.G.: data collection
- J.G., D.M., C.P., P.T., and N.R.: interpret results
- P.T. and N.R.: design and lead research
- All authors: writing manuscript

## Acknowledgements

J.G. acknowledges the support of Comisión Académica de Posgrado (CAP), CSIC Iniciación and PEDECIBA. P.T. and N.R. also acknowledges the support of PEDECIBA.

## Supplementary Material

**Figure S1:**
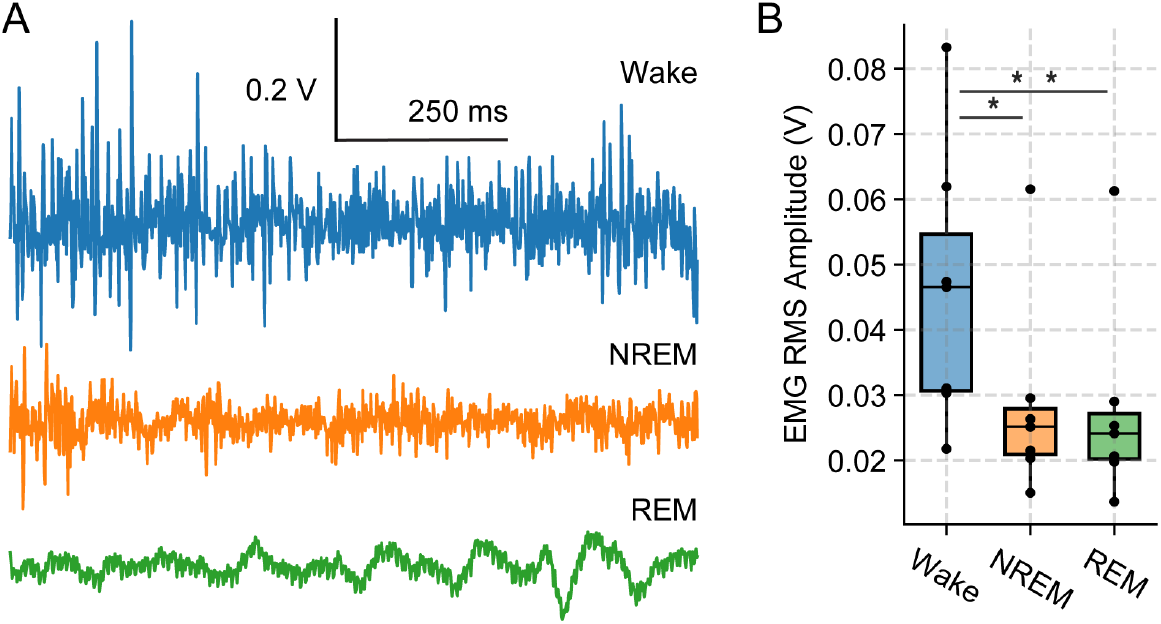
EMG variations for wakefulness (Wake), rapid-eye movement (REM) and non-REM (NREM) sleep. **A)** Electromyogram (EMG) traces for the different sleep-wake states. **B** Boxplots showing the root mean square (RMS) amplitude of the EMG activity for the each sleep state. * = *p* < 0.05 and ** = *p* < 0.01

**Figure S2:**
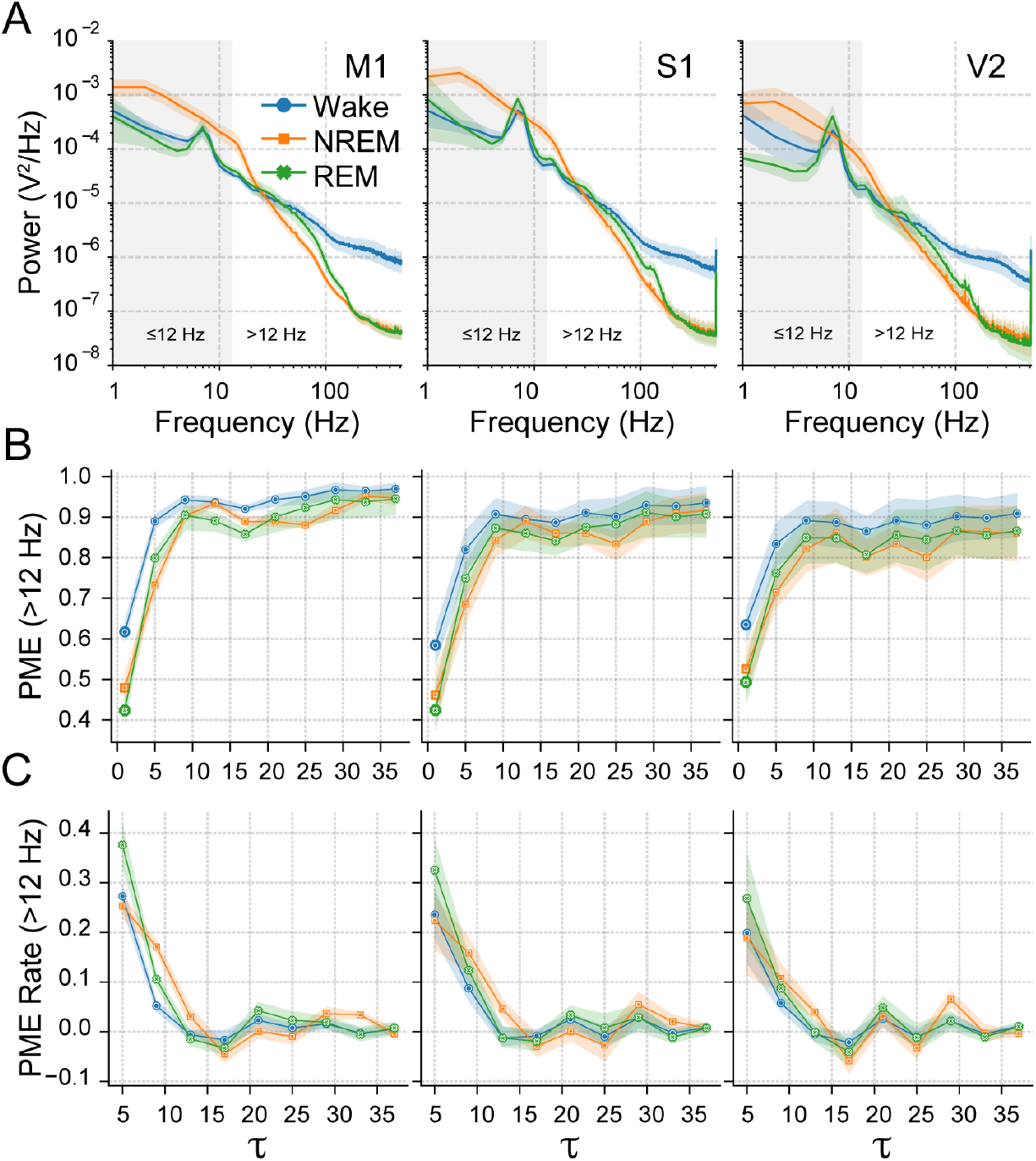
Power spectra and entropy variations for wakefulness (Wake), rapid-eye movement (REM) and non-REM (NREM) sleep. **A)** Power spectral densities for different cortical locations and sleep-wake states. Shaded rectangular areas signal the low frequency band (> 12 *Hz*). **B)** Permutation Minimum-Entropy (PME) values for the EEG’s high frequency-band as the embedding delay, *τ*, of the ordinal pattern (OP) is increased (i.e., data sampled at increasing steps) in each cortical location from Fig. 1A. **C)** Corresponding PME rates [Eq. (4)] from panel **B**, i.e., PME tangents as a function of *τ*. OP dimension in panels **B** and **C** is *D* = 5 data points (as in Fig. 1C). All panels show population averages (solid curves) with their 95% confidence interval (coloured shading) after 1000 bootstrap samples.

## References

Abásolo, D., Simons, S., Morgado da Silva, R., Tononi, G., and Vyazovskiy, V. V. (2015). Lempel-Ziv complexity of cortical activity during sleep and waking in rats. J Neurophysiol, 113(7):2742–2752.

Bandt, C. (2017). A new kind of permutation entropy used to classify sleep stages from invisible eeg microstructure. Entropy, 19(5).

Bandt, C. and Pompe, B. (2002). Permutation entropy: a natural complexity measure for time series. Physical review letters, 88(17):174102.

Bastos, A. M., Lundqvist, M., Waite, A. S., Kopell, N., and Miller, E. K. (2020). Layer and rhythm specificity for predictive routing. Proc Natl Acad Sci U S A, 117(49):31459–31469.

Bastos, A. M., Vezoli, J., Bosman, C. A., Schoffelen, J. M., Oostenveld, R., Dowdall, J. R., De Weerd, P., Kennedy, H., and Fries, P. (2015). Visual areas exert feedforward and feedback influences through distinct frequency channels. Neuron, 85(2):390–401.

Buzsáki, G., Anastassiou, C. A., and Koch, C. (2012). The origin of extracellular fields and currents–EEG, ECoG, LFP and spikes. Nat Rev Neurosci, 13(6):407–420.

Buzsáki, G. and Draguhn, A. (2004). Neuronal oscillations in cortical networks. Science, 304(5679):1926–1929.

Carr, M. F., Karlsson, M. P., and Frank, L. M. (2012). Transient slow gamma synchrony underlies hippocampal memory replay. Neuron, 75(4):700–713.

Cavelli, M., Castro-Zaballa, S., Mondino, A., Gonzalez, J., Falconi, A., and Torterolo, P. (2017). Absence of eeg gamma coherence in a local activated cortical state: a conserved trait of rem sleep. Translational Brain Rhythmicity, 21132017.

Cavelli, M., Rojas-Líbano, D., Schwarzkopf, N., Castro-Zaballa, S., Gonzalez, J., Mondino, A., Santana, N., Benedetto, L., Falconi, A., and Torterolo, P. (2018). Power and coherence of cortical high-frequency oscillations during wakefulness and sleep. European Journal of Neuroscience, 48(8):2728–2737.

Chase, M. H. and Morales, F. R. (1983). Subthreshold excitatory activity and motoneuron discharge during REM periods of active sleep. Science, 221(4616):1195–1198.

Chase, M. H., Soja, P. J., and Morales, F. R. (1989). Evidence that glycine mediates the postsynaptic potentials that inhibit lumbar motoneurons during the atonia of active sleep. J Neurosci, 9(3):743–751.

D’Andola, M., Rebollo, B., Casali, A. G., Weinert, J. F., Pigorini, A., Villa, R., Massimini, M., and Sanchez-Vives, M. V. (2018). Bistability, Causality, and Complexity in Cortical Networks: An In Vitro Perturbational Study. Cereb Cortex, 28(7):2233–2242.

Dasilva, M., Camassa, A., Navarro-Guzman, A., Pazienti, A., Perez-Mendez, L., Zamora-López, G., Mattia, M., and Sanchez-Vives, M. V. (2021). Modulation of cortical slow oscillations and complexity across anesthesia levels. Neuroimage, 224:117415.

Eichenlaub, J. B., Biswal, S., Peled, N., Rivilis, N., Golby, A. J., Lee, J. W., Westover, M. B., Halgren, E., and Cash, S. S. (2020). Reactivation of Motor-Related Gamma Activity in Human NREM Sleep. Front Neurosci, 14:449.

Evarts, E. V. (1964). Temporal patterns of discharge of pyramidal tract neurons during sleep and waking in the monkey. Journal of neurophysiology, 27(2):152–171.

Fagerholm, E. D., Scott, G., Shew, W. L., Song, C., Leech, R., Knöpfel, T., and Sharp, D. J. (2016). Cortical Entropy, Mutual Information and Scale-Free Dynamics in Waking Mice. Cereb Cortex, 26(10):3945–3952.

Gervasoni, D., Lin, S. C., Ribeiro, S., Soares, E. S., Pantoja, J., and Nicolelis, M. A. (2004). Global forebrain dynamics predict rat behavioral states and their transitions. J Neurosci, 24(49):11137–11147.

González, J., Cavelli, M., Tort, A. B., Torterolo, P., and Rubido, N. (2021). Off-periods reduce the complexity of neocortical activity during sleep. bioRxiv.

González, J., Cavelli, M., Mondino, A., Pascovich, C., Castro-Zaballa, S., Torterolo, P., and Rubido, N. (2019). Decreased electrocortical temporal complexity distinguishes sleep from wakefulness. Sci Rep, 9(1):18457.

González, J., Cavelli, M., Mondino, A., Pascovich, C., Castro-Zaballa, S., Torterolo, P., and Rubido, N. (2020). Electrocortical temporal complexity during wakefulness and sleep: an updated account. Sleep Science.

Hou, F., Zhang, L., Qin, B., Gaggioni, G., Liu, X., and Vandewalle, G. (2021). Changes in EEG permutation entropy in the evening and in the transition from wake to sleep. Sleep, 44(4).

Jordan, D., Stockmanns, G., Kochs, E. F., Pilge, S., and Schneider, G. (2008). Electroencephalographic order pattern analysis for the separation of consciousness and unconsciousness: an analysis of approximate entropy, permutation entropy, recurrence rate, and phase coupling of order recurrence plots. Anesthesiology, 109(6):1014–1022.

Kanayama, N., Sato, A., and Ohira, H. (2007). Crossmodal effect with rubber hand illusion and gamma-band activity. Psychophysiology, 44(3):392–402.

Kisley, M. A. and Cornwell, Z. M. (2006). Gamma and beta neural activity evoked during a sensory gating paradigm: effects of auditory, somatosensory and crossmodal stimulation. Clinical neurophysiology, 117(11):2549–2563.

Mateos, D. M., Gómez-Ramírez, J., and Rosso, O. A. (2021). Using time causal quantifiers to characterize sleep stages. Chaos, Solitons & Fractals, 146:110798.

Mondino, A., Hambrecht-Wiedbusch, V. S., Li, D., York, A. K., Pal, D., González, J., Torterolo, P., Mashour, G. A., and Vanini, G. (2021). Glutamatergic Neurons in the Preoptic Hypothalamus Promote Wakefulness, Destabilize NREM Sleep, Suppress REM Sleep, and Regulate Cortical Dynamics. J Neurosci, 41(15):3462–3478.

Nicolaou, N. and Georgiou, J. (2011). The use of permutation entropy to characterize sleep electroencephalograms. Clin EEG Neurosci, 42(1):24–28.

Nir, Y., Staba, R. J., Andrillon, T., Vyazovskiy, V. V., Cirelli, C., Fried, I., and Tononi, G. (2011). Regional slow waves and spindles in human sleep. Neuron, 70(1):153–169.

Oizumi, M., Albantakis, L., and Tononi, G. (2014). From the phenomenology to the mechanisms of consciousness: Integrated Information Theory 3.0. PLoS Comput Biol, 10(5):e1003588.

Ouyang, G., Dang, C., Richards, D. A., and Li, X. (2010). Ordinal pattern based similarity analysis for EEG recordings. Clin Neurophysiol, 121(5):694–703.

Pascovich, C., Castro-Zaballa, S., Mediano, P. A., Bor, D., Canales-Johnson, A., Torterolo, P., and Bekinschtein, T. A. (2021). Ketamine and sleep modulate neural complexity dynamics in cats. bioRxiv.

Pigorini, A., Sarasso, S., Proserpio, P., Szymanski, C., Arnulfo, G., Casarotto, S., Fecchio, M., Rosanova, M., Mariotti, M., Lo Russo, G., Palva, J. M., Nobili, L., and Massimini, M. (2015). Bistability breaks-off deterministic responses to intracortical stimulation during non-REM sleep. Neuroimage, 112:105–113.

Rényi, A. (1965). On the foundations of information theory. Revue de l’Institut International de Statistique, pages 1–14.

Rényi, A. et al. (1961). On measures of entropy and information. In Proceedings of the Fourth Berkeley Symposium on Mathematical Statistics and Probability, Volume 1: Contributions to the Theory of Statistics. The Regents of the University of California.

Richter, C. G., Thompson, W. H., Bosman, C. A., and Fries, P. (2017). Top-Down Beta Enhances Bottom-Up Gamma. J Neurosci, 37(28):6698–6711.

Rosanova, M., Fecchio, M., Casarotto, S., Sarasso, S., Casali, A. G., Pigorini, A., Comanducci, A., Seregni, F., Devalle, G., Citerio, G., Bodart, O., Boly, M., Gosseries, O., Laureys, S., and Massimini, M. (2018). Sleep-like cortical OFF-periods disrupt causality and complexity in the brain of unresponsive wakefulness syndrome patients. Nat Commun, 9(1):4427.

Sarasso, S., Boly, M., Napolitani, M., Gosseries, O., Charland-Verville, V., Casarotto, S., Rosanova, M., Casali, A. G., Brichant, J. F., Boveroux, P., Rex, S., Tononi, G., Laureys, S., and Massimini, M. (2015). Consciousness and Complexity during Unresponsiveness Induced by Propofol, Xenon, and Ketamine. Curr Biol, 25(23):3099–3105.

Sarasso, S., Casali, A. G., Casarotto, S., Rosanova, M., Sinigaglia, C., and Massimini, M. (2021). Consciousness and complexity: a consilience of evidence. Neuroscience of Consciousness. niab023.

Schartner, M. M., Pigorini, A., Gibbs, S. A., Arnulfo, G., Sarasso, S., Barnett, L., Nobili, L., Massimini, M., Seth, A. K., and Barrett, A. B. (2017). Global and local complexity of intracranial EEG decreases during NREM sleep. Neurosci Conscious, 2017(1):niw022.

Shannon, C. E. (1948). A mathematical theory of communication. The Bell system technical journal, 27(3):379–423.

Sitt, J. D., King, J. R., El Karoui, I., Rohaut, B., Faugeras, F., Gramfort, A., Cohen, L., Sigman, M., Dehaene, S., and Naccache, L. (2014). BrainLarge scale screening of neural signatures of consciousness in patients in a vegetative or minimally conscious state. Brain, 137(Pt 8):2258–2270.

Thul, A., Lechinger, J., Donis, J., Michitsch, G., Pichler, G., Kochs, E. F., Jordan, D., Ilg, R., and Schabus, M. (2016). EEG entropy measures indicate decrease of cortical information processing in Disorders of Consciousness. Clin Neurophysiol, 127(2):1419–1427.

Todorova, R. and Zugaro, M. (2019). ScienceIsolated cortical computations during delta waves support memory consolidation. Science, 366(6463):377–381.

Tononi, G. and Edelman, G. M. (1998). Consciousness and complexity. Science, 282(5395):1846–1851.

Valderrama, M., Crépon, B., Botella-Soler, V., Martinerie, J., Hasboun, D., Alvarado-Rojas, C., Baulac, M., Adam, C., Navarro, V., and Le Van Quyen, M. (2012). Human gamma oscillations during slow wave sleep. PLoS One, 7(4):e33477.

Varley, T. F., Denny, V., Sporns, O., and Patania, A. (2021). Topological analysis of differential effects of ketamine and propofol anaesthesia on brain dynamics. R Soc Open Sci, 8(6):201971.

Varley, T. F., Sporns, O., Puce, A., and Beggs, J. (2020). Differential effects of propofol and ketamine on critical brain dynamics. PLoS Comput Biol, 16(12):e1008418.

Vyazovskiy, V. V., Olcese, U., Lazimy, Y. M., Faraguna, U., Esser, S. K., Williams, J. C., Cirelli, C., and Tononi, G. (2009). Cortical firing and sleep homeostasis. Neuron, 63(6):865–878.

Watson, B. O., Levenstein, D., Greene, J. P., Gelinas, J. N., and Buzsáki, G. (2016a). Multi-unit spiking activity recorded from rat frontal cortex (brain regions mPFC, OFC, ACC, and M2) during wake-sleep episode wherein at least 7 minutes of wake are followed by 20 minutes of sleep. CRCNS.org.

Watson, B. O., Levenstein, D., Greene, J. P., Gelinas, J. N., and Buzsáki, G. (2016b). Network homeostasis and state dynamics of neocortical sleep. Neuron, 90(4):839–852.

Whitham, E. M., Pope, K. J., Fitzgibbon, S. P., Lewis, T., Clark, C. R., Loveless, S., Broberg, M., Wallace, A., DeLosAngeles, D., Lillie, P., Hardy, A., Fronsko, R., Pulbrook, A., and Willoughby, J. O. (2007). Scalp electrical recording during paralysis: quantitative evidence that EEG frequencies above 20 Hz are contaminated by EMG. Clin Neurophysiol, 118(8):1877–1888.

Wiesman, A. I., Koshy, S. M., Heinrichs-Graham, E., and Wilson, T. W. (2020). Beta and gamma oscillations index cognitive interference effects across a distributed motor network. Neuroimage, 213:116747.

Zanin, M., Zunino, L., Rosso, O. A., and Papo, D. (2012). Permutation entropy and its main biomedical and econophysics applications: a review. Entropy, 14(8):1553–1577.

Zunino, L., Olivares, F., and Rosso, O. A. (2015). Permutation min-entropy: An improved quantifier for unveiling subtle temporal correlations. EPL (Europhysics Letters), 109(1):10005.

